# Csf1 from marrow adipogenic precursors is required for osteoclast formation and hematopoiesis in bone

**DOI:** 10.1101/2022.07.27.501742

**Authors:** Leilei Zhong, Jiawei Lu, Jiankang Fang, Lutian Yao, Wei Yu, Tao Gui, Nicholas Holdreith, Catherine Bautista, Yongwon Choi, Jean X. Jiang, Shuying Yang, Wei Tong, Nathaniel Dyment, Ling Qin

## Abstract

Colony stimulating factor 1 (Csf1) is an essential growth factor for osteoclast progenitors and thus an important regulator for bone resorption. It remains elusive which mesenchymal cells synthesize *Csf1* stimulating osteoclastogenesis. We recently identified a novel mesenchymal cell population, marrow adipogenic lineage precursors (MALPs), in bone. Single cell RNA-sequencing indicated specific expression of *Csf1* in MALPs, which is further increased during aging. To investigate its role, we constructed *Csf1 CKO* mice using *Adipoq-Cre*. These mice showed increased femoral trabecular bone over time, but their cortical bone appeared normal. In comparison, depletion of Csf1 in the entire mesenchymal lineage using *Prx1-Cre* led to a more striking high bone mass phenotype, suggesting that additional mesenchymal subpopulations secrete Csf1. TRAP staining revealed diminished osteoclasts in the femoral secondary spongiosa region of *Csf1 CKO*^*Adipoq*^ mice, but not at the chondral-osseous junction nor at the endosteal surface of cortical bone. Moreover, *Csf1 CKO*^*Adipoq*^ mice were resistant to LPS-induced calvarial osteolysis. Bone marrow cellularity, hematopoietic progenitors, and macrophages were also reduced in these mice. Taken together, our studies demonstrate that MALPs are a critical player in controlling bone remodeling and hematopoiesis.

## Introduction

Bone is maintained by a fine balance between bone formation by osteoblasts and bone resorption by osteoclasts. These two types of functional cells originate from different lineages of stem cells, with the former one derived from bone marrow mesenchymal stem cells (MSCs) and the latter one from hematopoietic stem cells (HSCs). In addition to osteoblasts and osteoclasts, MSCs and HSCs also give rise to many other cell subpopulations that co-exist inside bone (1, 2). Crosstalk among mesenchymal and hematopoietic subpopulations plays a critical role in bone homeostasis, and disruption of it shifts the balance of bone remodeling, leading to bone disorders (3).

As a specific type of macrophage, osteoclasts are differentiated from monocytes in the myeloid subpopulation of hematopoietic lineage cells (4). Colony stimulating factor 1 (Csf1), also known as macrophage colony-stimulating factor (M-Csf), is essential for osteoclastogenesis due to its actions in promoting proliferation, survival, and differentiation of monocytes and macrophages (5). The expression of its receptor, Csf1r, is low in immature myeloid precursor cells and increases as the myeloid cells mature (6). Receptor activator of NF-κB ligand (RANKL) is another indispensable cytokine for promoting osteoclastogenesis (7-9). Csf1/Csf1r signaling induces the expression of the receptor activator of NF-κB (RANK), a receptor for RANKL, to further facilitate the formation of mature osteoclasts (10). Through alternative splicing, Csf1 can be expressed into a secreted glycoprotein, a secreted proteoglycan, or a membrane-spanning cell surface glycoprotein (11). A spontaneous inactivating mutation in the mouse *Csf1* coding region causes a complete loss of Csf1 protein (12). The resultant homozygous *Csf1op/op* mice are osteopetrotic due to osteoclast deficiency, but this bone abnormality recovers over the first few months of life (13). Similarly, a spontaneous recessive mutation (*toothless, tl*) in the rat *Csf1* gene leads to a more complete loss of osteoclasts, resulting in more severe osteopetrosis than *Csf1op/op* mice with no improvement when rats age (14). Knocking out *Csf1r* gene in mice generates similar or even more severe skeletal phenotypes than *Csf1* deficient mice (15), further demonstrating the important role of Csf1/Csf1r signaling in controlling bone resorption.

Past studies have demonstrated the expression of Csf1 in a wide range of cells, such as fibroblasts, endothelial cells, keratinocytes, astrocytes, myoblasts, breast and uterine epithelial cells (16). Primary cell culture experiments indicated that osteoblasts synthesize both soluble and membrane bound forms of Csf1 (17). Interestingly, targeting expression of soluble Csf1 by osteoblast-specific promoter (osteocalcin) rescues the osteopetrotic bone defect in *Csf1op/op* mice (18). Csf1 is also expressed in osteocytes (19). Mice with osteocyte-specific *Csf1* deficiency were generated using *Dmp1-Cre*. These mice showed increased trabecular bone mass in tibiae and vertebrates (19). Therefore, the prevalent view is that bone forming cells, including osteoblasts and osteocytes, are the major producers of Csf1 to control osteoclast formation.

In addition to bone forming cells, bone marrow MSCs also give rise to marrow adipocytes. With the advance of single cell transcriptomics technology, we and others recently dissected the heterogeneity of bone marrow mesenchymal lineage cells and delineated the bi-lineage differentiation trajectories of MSCs (20-23). Specifically, we identified a novel mesenchymal subpopulation termed “marrow adipogenic lineage precursors” (MALPs) that expresses many adipocyte markers but contains no lipid droplets (24). As precursors for lipid-laden adipocytes (LiLAs), MALPs exist abundantly as pericytes and stromal cells, forming an extensive 3D network inside the marrow cavity. Interestingly, computational analysis revealed MALPs to be the most interactive mesenchymal cells with monocyte-macrophage lineage cells due to their high and specific expression of several osteoclast regulatory factors, including RANKL and Csf1 (25). Studies from our group and others have demonstrated that MALP-derived RANKL is critical for promoting bone resorption during physiological and pathological conditions (25, 26). Since Csf1 is also important for osteoclastogenesis, here we generated *Csf1* conditional knockout mice using *Adipoq-Cre* driver and examine the role of MALP-derived Csf1 in bone homeostasis and diseases. Our findings identified MALPs as a major source of Csf1 in bone and reinforce our discovery that MALPs is a novel key player in controlling bone resorption. Furthermore, we found that MALP-derived Csf1 contributes to hematopoiesis in the bone marrow, particularly macrophage production.

## Results

### ScRNA-seq data suggest MALPs as the major source of Csf1 in bone

To examine the cellular sources of Csf1, we integrated scRNA-seq datasets of bone marrow cells we previously generated from 1, 3, and 16-month-old mice (Fig. 1A) (23). These experiments were initially designed to analyze mesenchymal lineage cells in bone. Since they also included hematopoietic, endothelial, and mural cells, those scRNA-seq datasets indeed represent the overall cellular composition of bone tissue. Interestingly, we found that in young and adult mice, the major Csf1 producing cells are MALPs, followed by lineage committed progenitors (LCPs) and early mesenchymal progenitors (EMPs, Fig. 1B). Aging not only increased *Csf1* expression in these cells, but also initiated its expression in some other subpopulations, such as mesenchymal cells [early mesenchymal progenitors (EMPs), late mesenchymal progenitors (LMPs), and osteoblasts], hematopoietic cells (macrophages, granulocyte progenitors, and red blood cells), endothelial cells, and mural cells. In aged mice, MALPs were markedly expanded while EMPs and LMPs were shrunk in numbers (Fig. 1C).

**Figure 1.**
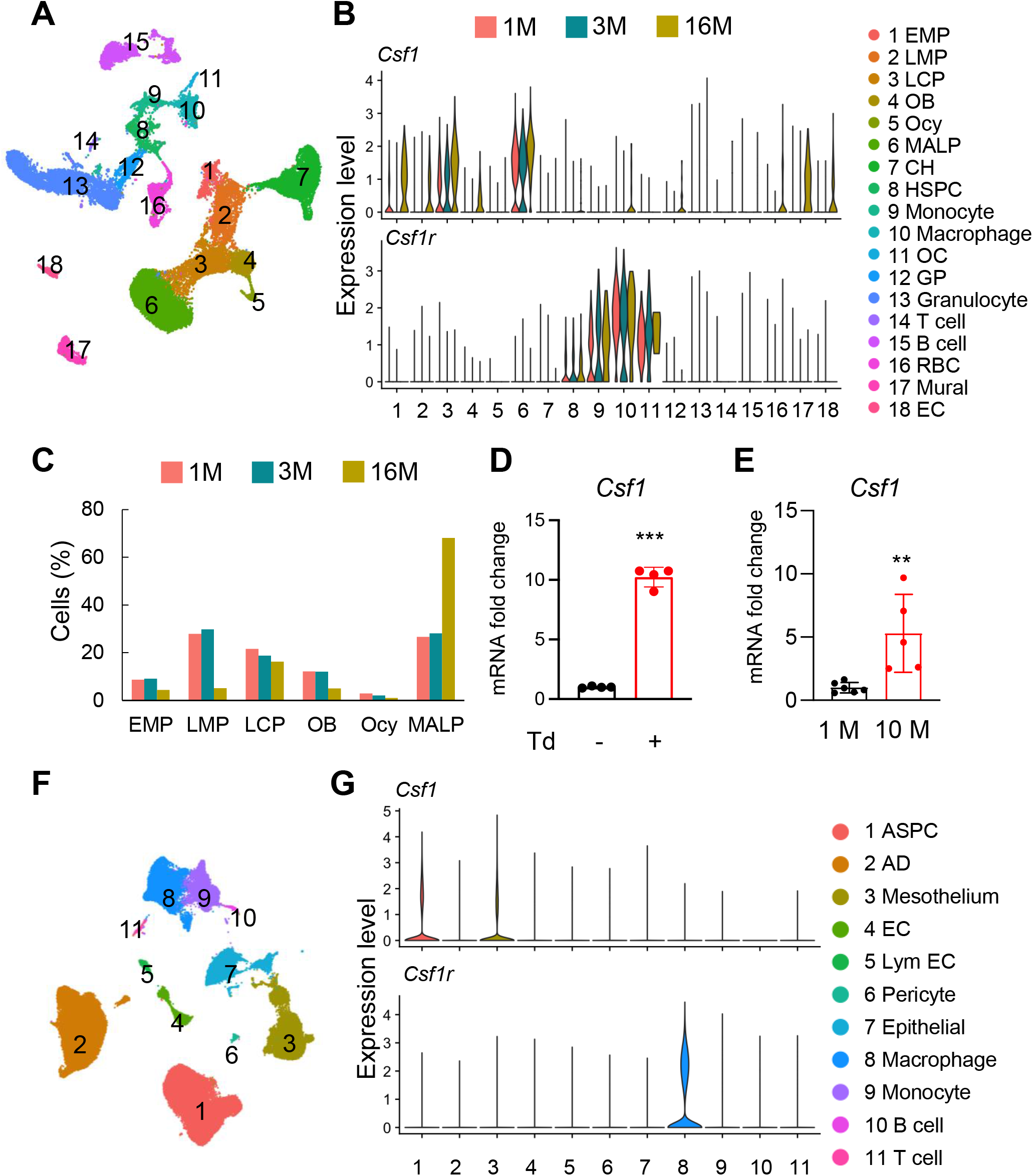
*Csf1* expression in bone is mainly contributed by MALPs in an age-dependent manner. (A) The integrated scRNA-seq datasets of sorted bone marrow Td+ cells from 1, 3, and 16- month-old *Col2/Td* mice (n=11 mice). The UMAP plot is presented to show cell clustering. (B) Violin plots of *Csf1* and its receptor *Csf1r* in bone marrow cells at different ages. EMP: early mesenchymal progenitor; LMP: late mesenchymal progenitor; LCP: lineage committed progenitor; OB: osteoblast; Ocy: osteocyte; CH: chondrocyte; EC: endothelial cell; HSPC: hematopoietic stem and progenitor cell; OC: osteoclast; GP: granulocyte progenitor; RBC: red blood cell; EC: endothelial cell. (C) The percentages of cells in each cell cluster within bone marrow mesenchymal lineage cells are quantified based on UMAP distribution. (D) qRT-PCR analysis of *Csf1* expression in sorted Td+ and Td- bone marrow cells from Adipoq/Td mice at 3 months of age. ***, p<0.001 Td+ vs Td- cells. (E) qRT-PCR analysis of *Csf1* expression in young and aged bone marrow from *WT* mice. **, p<0.01 10 M vs 1 M mice. (F) SnRNA-seq analysis of inguinal and perigonadal adipose tissues from 16-week-old mice. The UMAP plot is presented to show cell clustering. (G) Violin plots of Csf1 and its receptor *Csf1r* in individual cell subpopulation from adipose tissues. AD: adipocytes; EC: endothelial cells; Lym EC: lymphatic endothelial cells.

Thus, our scRNA-seq data predict an increased amount of Csf1 in mouse bone marrow during aging. Concomitantly, *Csf1r* was mainly expressed in monocytic subpopulations, such as monocytes, macrophages, and osteoclasts, in an age-independent manner (Fig. 1B). In line with previous reports (27, 28), HSCs also expressed *Csf1r* at a low level. To validate these sequencing data, we sorted Td+ cells from the bone marrow of 3-month-old *Adipoq-Cre Rosa-tdTomato* (*Adipoq/Td*) mice, a MALP reporter line. While Td+ cells only make up ∼0.5% of bone marrow cells (23), qRT-PCR analysis revealed that their expression of *Csf1* is 10.2-fold higher than Td- cells (Fig. 1D). Furthermore, bone marrow cells from 10-month-old mice expressed 5.9-fold more *Csf1* than those from 1-month-old mice (Fig. 1E).

MALPs can further differentiate into LiLAs (23). It is technically challenging to isolate mouse bone marrow LiLAs and examine their gene expression profile. To understand whether LiLAs express *Csf1*, we analyzed a single nucleus RNA-sequencing (snRNA-seq) dataset from white adipose tissue of 16-week-old mice that contains LiLAs (GSE number 176171, Fig. 1F) (29). Surprisingly, LiLAs did not express *Csf1* at a detectable level. Instead, adipose stem and progenitor cells (ASPCs) and mesothelial cells were the major Csf1 producing cells, albeit their expression levels were much lower compared to MALPs in the bone marrow (Fig. 1G). As expected, the highest *Csf1r* expression was detected in adipose tissue resident macrophages (Fig. 1G). Taken together, our scRNA-seq data indicate that MAPLs are the major Csf1 producing cells in bone tissue.

### Mice with Csf1 deficiency in MALPs have high trabecular bone mass

Our previous studies demonstrate that adipocyte-specific *Adipoq-Cre* targets MALPs and LiLAs, but not osteoblasts and osteocytes, in young and adult mice (23, 25). Thus, we constructed *Adipoq-Cre Csf1*^*flox/flox*^ *(Csf1 CKO*^*Adipoq*^) mice to investigate the role of MALPs-derived Csf1. These mice were born at the expected frequency. They grew normally with comparable body weight, femoral length, and tooth eruption as their *WT* siblings (Fig. 2A-C). Their peripheral white adipose tissue also appeared normal, with similar adipocyte size, number and vasculature as *WT* mice (Fig. S1). The *Csf1* mRNA level in *CKO* bone marrow was reduced to 43% of *WT* bone marrow (Fig. 2D). This result not only validates the efficiency of the knockout but also supports MALPs as the major source of Csf1 in the bone marrow. On the contrary, Csf1 expression in the cortical bone was not changed (Fig. 2D), suggesting that osteocytes are not affected in *Csf1 CKO*^*Adipoq*^ mice.

**Figure 2.**
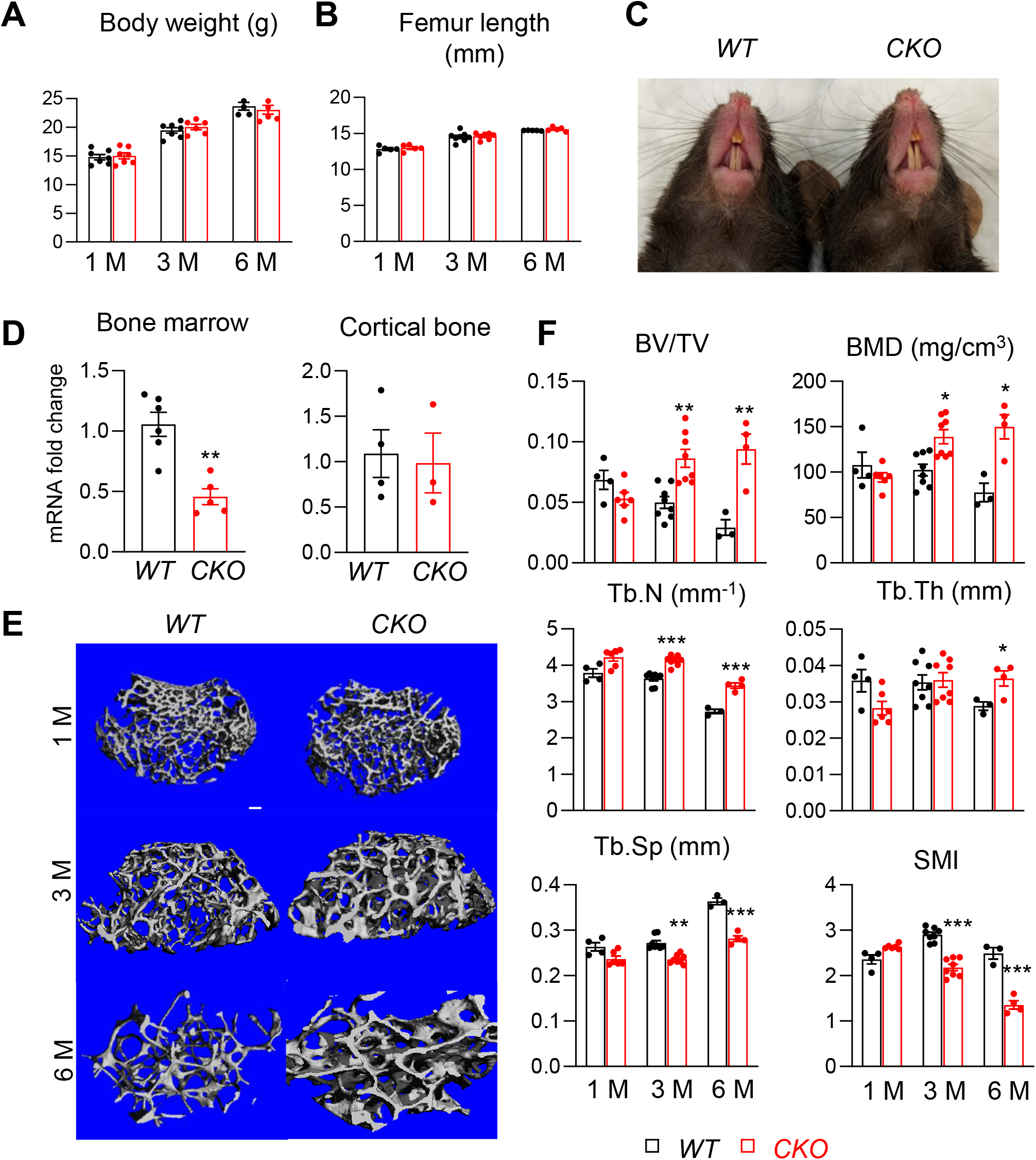
*Csf1 CKO*^*Adipoq*^ mice have high trabecular bone mass in long bone. (A, B) *Csf1 CKO*^*Adipoq*^ mice have normal body weight (A) and femoral length (B) at 1, 3, and 6 months of age. n=5-7 mice/group. (C) Tooth eruption is also normal in *CKO* mice at 1 month of age. (D) qRT-PCR analysis of *Csf1* mRNA in bone marrow and cortical bone of *WT* and *Csf1 CKO*^*Adipoq*^ mice at 3 months of age. n=3-6 mice/group. (E) 3D microCT reconstruction of femoral secondary spongiosa region from 1, 3 and 6 month- old mice reveals a drastic increase of trabecular bone in female *Csf1 CKO*^*Adipoq*^ mice compared to *WT* mice. Scale bar=100 µm. (F) MicroCT measurement of trabecular bone structural parameters. BV/TV: bone volume fraction; BMD: bone mineral density; Tb.N: trabecular number; Tb.Th: trabecular thickness; Tb.Sp: trabecular separation; SMI: structural model index. n=3-8 mice/group. *, p<0.05; **, p,0.01; ***, p<0.001 *CKO* vs *WT*.

MicroCT analysis of femoral trabecular bone revealed substantial osteopetrosis in these *CKO* mice starting from 3 months of age (Fig. 2E, F). Compared to *WT* siblings, *CKO* mice showed significantly elevated bone volume fraction (BV/TV, 1.7-fold) and bone mineral density (BMD, 1.4-fold) at 3 months of age and more enhancement (3.2- and 1.9-fold, respectively) at 6 months of age. These changes were accompanied by increased trabecular number (Tb.N, 14% and 26% at 3 and 6 months of age, respectively) and trabecular thickness (Tb.Th, 26% at 6 months of age), and decreased trabecular separation (Tb.Sp, 13% and 22% at 3 and 6 months of age, respectively). The reduced structure model index (SMI) confirmed the improved trabecular bone microarchitecture in *CKO* mice. Meanwhile, femoral cortical bone structure was not altered (Fig. S2). To our surprise, while MALPs are present in vertebral bone marrow (23), we did not detect any structural changes in vertebral trabecular bone (Fig. S3).

### Csf1 from other mesenchymal lineage cells regulate bone structure

To explore whether Csf1 from MALPs plays a dominant role in regulating bone structure, we generated *Prx1-Cre Csf1*^*flox/flox*^ *(Csf1 CKO*^*Prx1*^) mice to knockout *Csf1* in all mesenchymal lineage cells. Like *Csf1 CKO*^*Adipoq*^ mice, these *Csf1 CKO*^*Prx1*^ mice had normal tooth eruption (data not shown). At 2-3 months of age, these mice showed more drastic skeletal abnormalities than *Csf1 CKO*^*Adipoq*^. Their femoral length was reduced by 13% (Fig. 3A), indicating a defect in growth plate development. Their trabecular BV/TV and BMD were remarkably increased by 10.3-fold and 5.7-fold, respectively, due to increased Tb.N (2.3-fold) and Tb.Th (2.7-fold), and decreased Tb.Sp (49%, Fig. 3B-D). Furthermore, their cortical bone was also enlarged. We observed significantly increased periosteal perimeter (Ps.Pm), endosteal perimeter (Ec.Pm), and cortical bone area (Ct.Ar) in *Csf1 CKO*^*Prx1*^ mice compared to *WT* mice (Fig. 3E, F). Interestingly, the endosteal surface in *Csf1 CKO*^*Prx1*^ mice was not as smooth as that in *WT* mice, due to the extensive trabecular bone remained at the midshaft area. Since *Prx1-Cre* labels the entire mesenchymal lineage cells in bone, including chondrocytes, mesenchymal progenitors, osteoblasts, and osteocytes, these data suggest that mesenchymal cells other than MALPs also contribute Csf1 to regulate bone structure.

**Figure 3.**
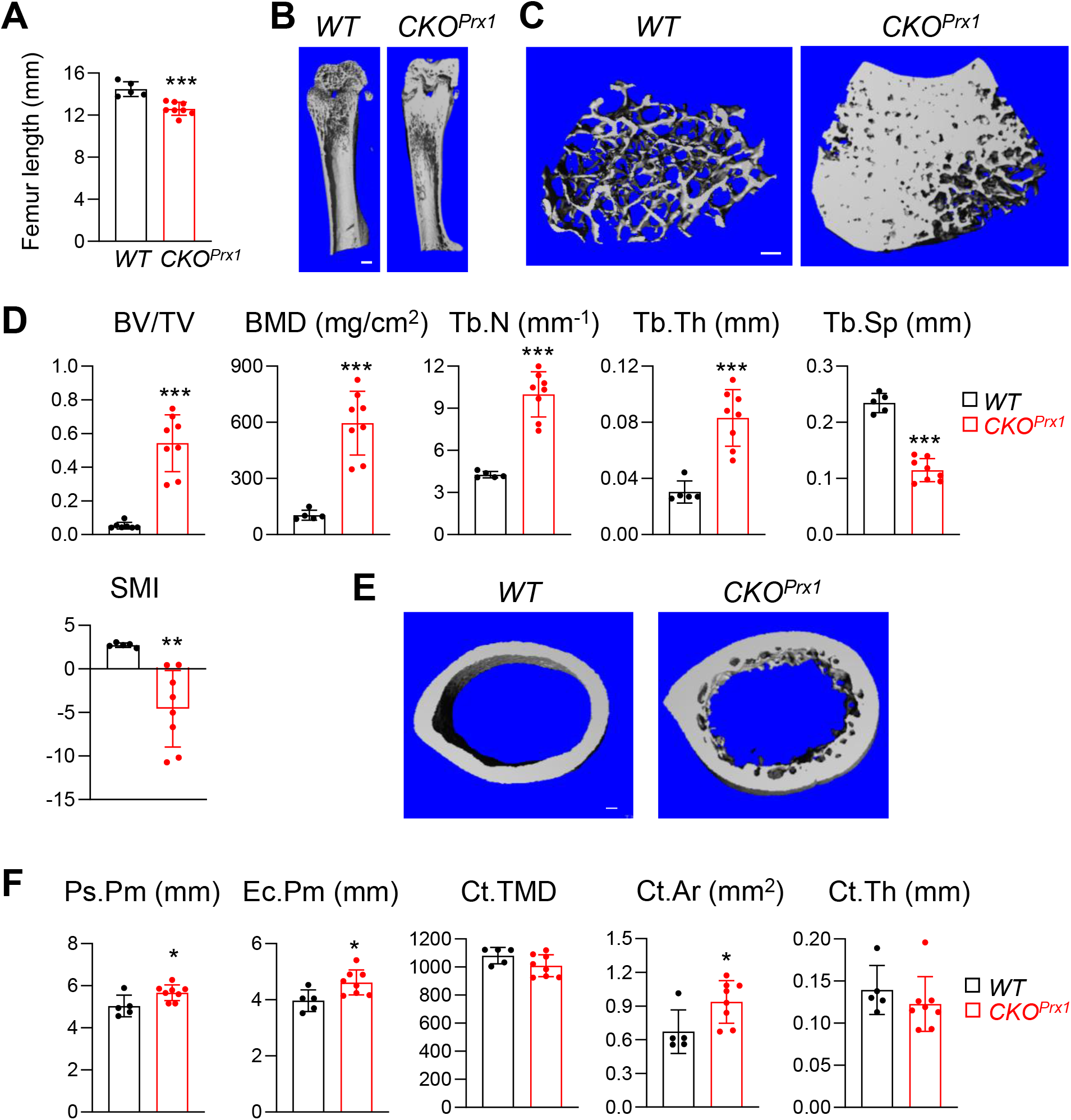
*Csf1* depletion in all mesenchymal cells using *Prx1-Cre* affects bone growth and causes severe osteopetrosis. (A) Femur length measurement in *Csf1 CKO*^*Prx1*^ mice at 2-3 months of age (n=5-8 mice/group). (B) 3D microCT reconstruction of whole femurs from *WT* and *Csf1-CKO*^*Prx1*^ mice. Scale bar=1 mm. (C) 3D microCT reconstruction of femoral secondary spongiosa region from *WT* and *Csf1- CKO*^*Prx1*^ mice. Scale bar=100 µm. (D) MicroCT measurement of trabecular bone structural parameters. BV/TV: bone volume fraction; BMD: bone mineral density; Tb.N: trabecular number; Tb.Th: trabecular thickness; Tb.Sp: trabecular separation; SMI: structural model index. n=5–8 mice/group. (E) 3D microCT reconstructions of femoral cortical bone. Scale bar=100 µm. (F) MicroCT measurement of cortical bone structural parameters. Ps.Pm: periosteal perimeter; Ec.Pm: endosteal perimeter; Ct.TMD: cortical tissue mineral density. Ct.Ar: cortical area; Ct.Th: cortical thickness. n=5–8 mice/group. *, p<0.05; **, p,0.01; ***, p<0.001 *CKO*^*Prx1*^ vs *WT*.

### Csf1deficiency in MALPs suppresses bone resorption but does not affect bone formation

To delineate the cellular mechanism of high bone mass in *Csf1 CKO*^*Adipoq*^ mice, we quantified osteoclasts at different locations of long bones (Fig. 4A, B). TRAP+ osteoclasts in the secondary spongiosa (SS) were remarkably reduced by 58%. However, no changes in osteoclasts were detected at the chondral-osseous junction (COJ) and the endosteal surface of cortical bone. To examine osteoclast progenitors in *Csf1 CKO*^*Adipoq*^ mice, we harvested bone marrow macrophages (BMMs) for in vitro osteoclastogenesis assay. Cells from both *WT* and *CKO* mice generated similar numbers of osteoclasts in culture (Fig. 4C, D), suggesting that the differentiation ability of osteoclast progenitors is maintained in *Csf1 CKO*^*Adipoq*^ mice.

**Figure 4.**
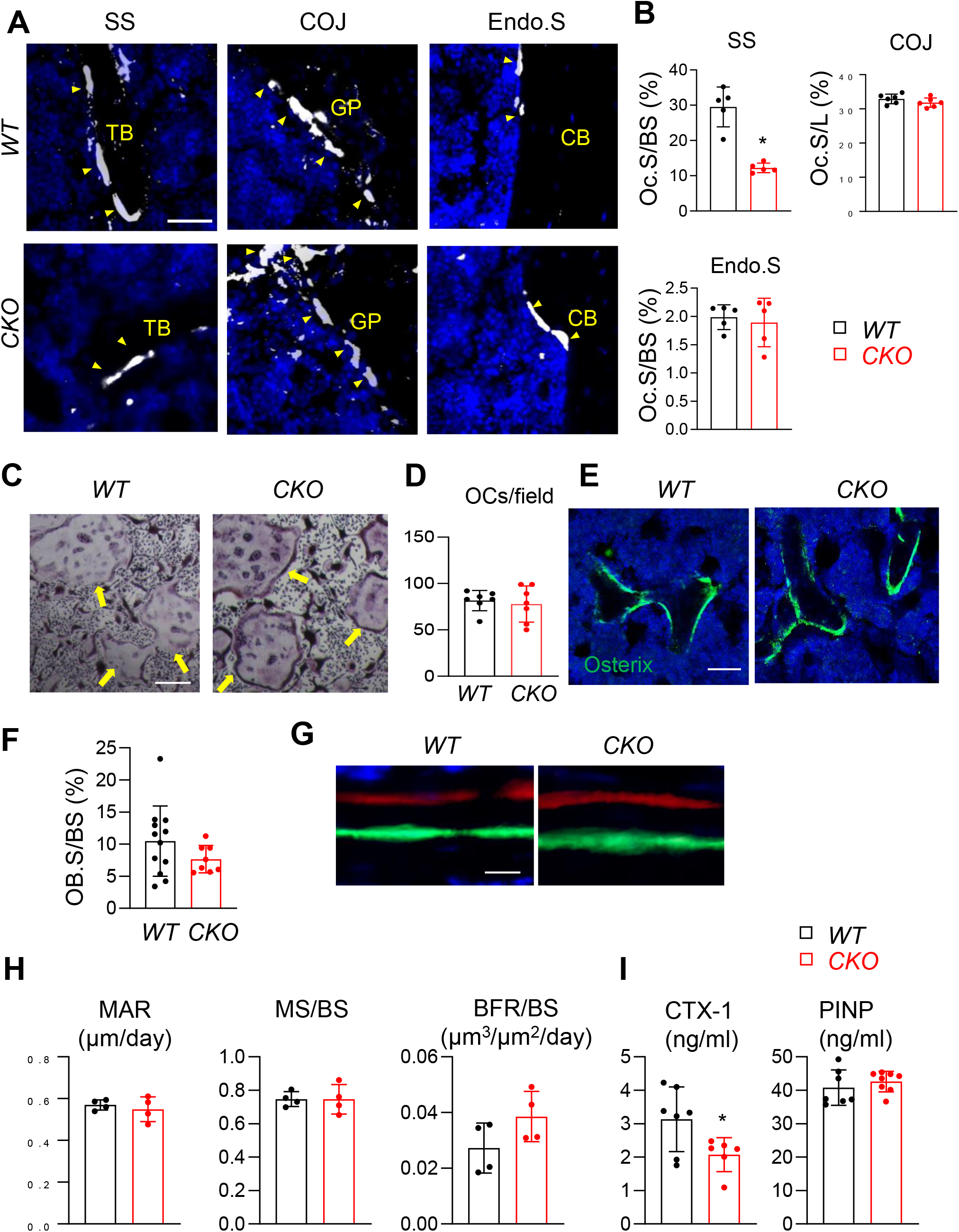
*Csf1* deletion in MALPs suppresses bone resorption but not bone formation. (A) Representative fluorescent TRAP staining images of femoral long bones from *WT* and *Csf1 CKO*^*Adipoq*^ mice at 3 months of age show TRAP^+^ osteoclasts at different skeletal sites: secondary spongiosa (SS), chondro-osseous junction (COJ), and endosteal surface (Endo.S). TB: trabecular bone; CB: cortical bone. Scale bar=50 μm. (B) Quantification of osteoclast surface (Oc.S) at three skeletal sites. BS: bone surface. L: COJ length. n=5 mice/group. *, p<0.05 *CKO* vs *WT*. (C) Representative TRAP staining images of osteoclast culture derived from *WT* and *Csf1 CKO*^*Adipoq*^ BMMs at 7 days after addition of RANKL and Csf1. Scale bar=200 μm. (D) Quantification of TRAP^+^ multinucleated cells (>3 nuclei/cell) per field. n=7 mice/group. (E) Representative Osterix staining of trabecular bone from *WT* and *Csf1 CKO*^*Adipoq*^ femurs. Scale bar=50 μm. (F) Quantification of osteoblast surface (OB.S). BS, bone surface. n=8-12 mice/group. (G) Representative double labeling of trabecular bone from *WT* and *Csf1 CKO*^*Adipoq*^ femurs. (H) Bone formation activity is quantified. MAR: mineral apposition rate; MS: mineralizing surface; BFR: bone formation rate. n=4 mice/group. (I) Serum ELISA analysis of bone resorption marker (CTX-1) and formation marker (PINP) in *WT* and *CKO* mice. n=6-8 mice/group. *, p<0.05 *CKO* vs *WT*.

We also measured bone formation activity in 3-month-old *Csf1 CKO*^*Adipoq*^ femurs. No significant difference was detected in Osterix+ osteoblasts at the trabecular bone surface in *WT* and *CKO* femurs (Fig. 4E, F). Dynamic bone histomorphometry showed a similar distance between the two fluorescent dyes injected at 9 and 2 days, respectively, in *WT* and *CKO* mice before euthanization (Fig. 4G). Quantification revealed comparable mineral apposition rate (MAR), mineralization surface (MS/BS), and bone formation rate (BFR/BS, Fig. 4H).

Furthermore, bone resorption marker (CTX-1) was significantly reduced in *CKO* serum, while bone formation marker (PINP) was not changed (Fig. 4I).

### Pathological bone loss is attenuated in Csf1 CKO^Adipoq^ mice

Lipopolysaccharide (LPS) injection above mouse calvaria induces osteolysis that mimics bacteria-induced bone loss. Since we previously identified the existence of MALPs in calvarial bone marrow (25), we applied this osteolysis model to *Csf1 CKO*^*Adipoq*^ mice. One week after LPS injection, we found a drastic increase in bone destruction in *WT* calvaria but very minor destruction in *Csf1 CKO*^*Adipoq*^ calvaria (Fig. 5A, B). TRAP staining revealed that LPS-induction of TRAP+ osteoclasts in calvaria is mostly attenuated in *CKO* mice (Fig. 5C, D). Taken together, our data showed that MALPs-derived Csf1 is important for osteoclastogenesis in diseased bones.

**Figure 5.**
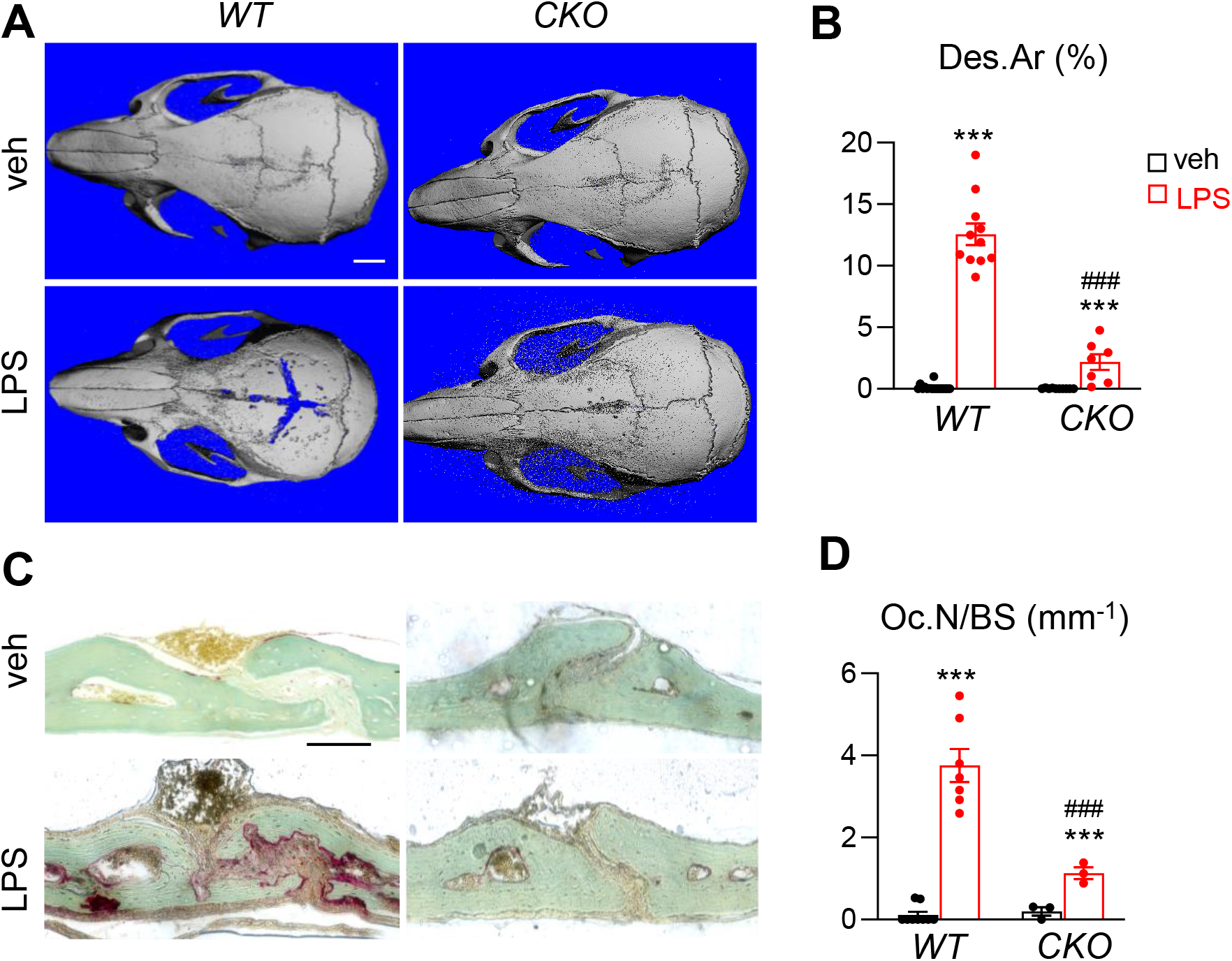
*Csf1 CKO*^*Adipoq*^ mice are protected from LPS-induced calvarial bone lesions. (A) Representative 3D microCT reconstruction of mouse calvaria after 1 week of vehicle (veh, PBS) or LPS injections. Scale bar=2 mm. (B) The percentage of bone destruction area (Des.Ar) in calvaria was quantified. n=7-16 mice/group. (C) Representative images of calvaria coronal section stained by TRAP. Scale bar=2 mm. (D) Quantification of osteoclast number (Oc.N) in calvaria. n=3-9 mice/group. ***, p<0.001 veh vs LPS; ###, p<0.001 *CKO* vs *WT*.

### Csf1 CKO^Adipoq^ mice have defective hematopoiesis in the bone marrow

Csf1 plays multifaceted roles in bone and blood tissues (16). We next investigated the role of MALPs-derived Csf1 in regulating hematopoiesis and bone marrow vasculature. Interestingly, bone marrow cellularity started to decline in *Csf1 CKO*^*Adipoq*^ mice at 3 months of age (18%) and further reduced at 6 months (43%, Fig. 6A), suggesting that hematopoiesis is suppressed.

**Figure 6.**
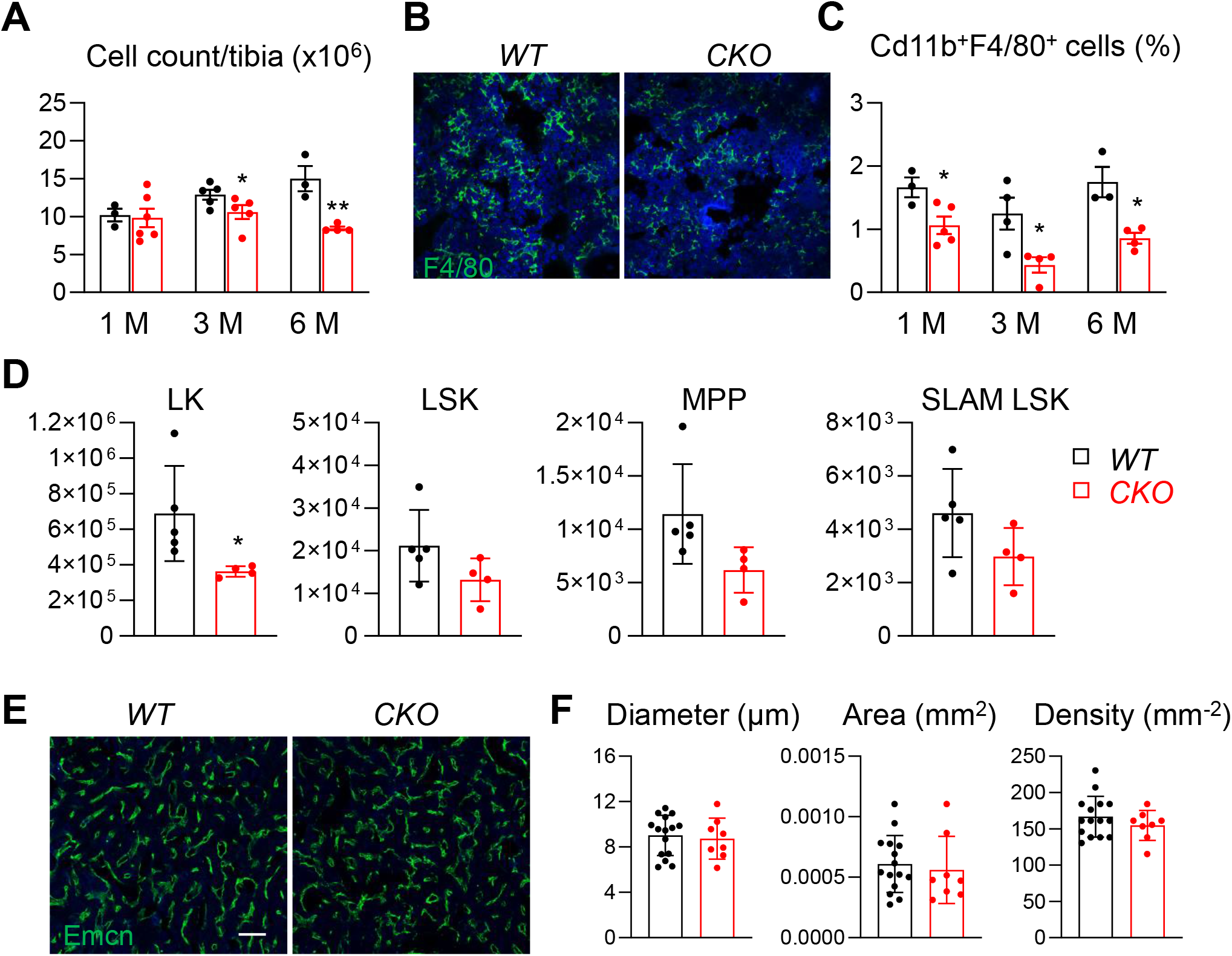
Bone marrow cellularity, hematopoietic progenitors, and macrophages are reduced in *Csf1 CKO*^*Adipoq*^ mice. (A) Bone marrow cellularity was quantified in *WT* and *Csf1 CKO*^*Adipoq*^ mice at 1, 3, and 6 months of age. *, p<0.05; **, p<0.01 *CKO* vs *WT*. (B) Representative fluorescent images of F4/80 staining in *WT* and *Csf1 CKO*^*Adipoq*^ bone marrow. (C) Flow analysis of Cd11+F4/80+ bone marrow macrophages at 1, 3, and 6 months of age. (D) Cell counts of hematopoietic stem and progenitor cells. n=4-5 mice/group. LK=Lineage- cKit^+^, LSK=Lineage^-^Sca1^+^cKit^+^, MPP=Lineage^-^Sca1^+^cKit^+^CD48^+^CD150^-^, SLAM LSK=Lineage-Sca1^+^cKit^+^CD48^-^CD150^+^. *, p<0.05 *CKO* vs *WT*. (E) Representative fluorescent images of bone marrow vasculature stained by Endomucin (Emcn). Scale bar=100 µm. (F) Quantification of bone marrow vessel diameter, density, and area. n=8-15 mice/group.

Macrophages are the major cell target of Csf1. Immunostaining revealed fewer F4/80+ macrophages in *CKO* bone marrow (Fig. 6B). Flow cytometric analysis confirmed that Cd11b+F4/80+ bone marrow macrophages were reduced by 36%, 65%, and 51%, in *CKO* mice compared to *WT* at 1, 3, and 6 months of age, respectively (Fig. 6C).

In addition to macrophages, we also found that hematopoietic stem and progenitors cells (HSPCs) are reduced in the bone marrow of *Csf1 CKO*^*Adipoq*^ mice (Fig. 6D). While only lineage-cKit^+^ (LK) cells displayed a significant decrease, more primitive progenitors, such as LSK, MPP, and SLAM LSK, showed a trend of reduction. However, these changes did not result in significant alterations in peripheral blood compositions (Fig. S4A) and spleen weight (Fig. S4B).

A previous study reported a reduction of bone marrow vasculature in the primary spongiosa area in mice with *Csf1* global deficiency (30). However, we did not detect obvious changes in marrow vasculature in *Csf1 CKO*^*Adipoq*^ mice (Fig. 6E, F), indicating that Csf1 from cells other than MALPs regulates angiogenesis. In summary, MALPs-derived Csf1 is required for regulating hematopoiesis, particularly macrophage production, in the bone marrow.

## Discussion

In bone, mesenchymal stem and progenitor cells undergo two divergent differentiation routes to produce osteogenic and adipogenic cells, respectively. In the past, bone research has mostly centered on osteogenic cells, including osteoblasts and osteocytes, because of their bone matrix synthesis ability, but paid less attention to marrow adipogenic cells. Recently, reports from our group and others discovered a group of marrow mesenchymal cells that express most adipocyte markers but do not contain lipid droplets (23, 31-33). Those cells are termed either “MALPs” based on their differentiation status or “MACs” (marrow Adipoq+ cells) based on their specific expression of Adipoq. Interestingly, our previous scRNA-seq study revealed that those adipogenic cells specifically and highly expressed two essential osteoclast regulatory factors, RANKL and Csf1 (23, 24). In our last report, we demonstrated that RANKL from MALPs promotes osteoclast formation and bone loss under normal and diseased conditions (25). In this study, we found that Csf1 from MALPs shares similar action in regulating bone resorption. Since all aspects of osteoclast formation and functions are regulated by RANKL and Csf1, our data firmly establish MALPs as one of the major bone cell types that control bone resorption.

In our previous study of mice with RANKL deficiency in MALPs (*Rankl CKO*^*Adipoq*^ mice), we proposed that osteoclast formation is controlled by various mesenchymal subpopulations in a site-dependent fashion. Our current research further substantiates this conclusion. We found that *Csf1 CKO*^*Adipoq*^ mice, similar to *Rankl CKO*^*Adipoq*^ mice, exhibit the trabecular bone phenotype but have normal long bone growth and cortical bone. However, *Csf1 CKO*^*Prx1*^ mice showed much more severe skeletal phenotypes. These mice have shortened long bones, in line with previous reports that global Csf1 deficiency in rodents is accompanied by progressive chondrodysplasia of the growth plate and altered endochondral ossification (30, 34). These data imply chondrocyte-derived Csf1 regulates cartilage-to-bone remodeling during endochondral ossification. *Csf1 CKO*^*Prx1*^ mice also have thickened cortical bones, suggesting a role of osteocytes in regulating cortical bone structure. Note that scRNA-seq data generated from our group and others do not contain hypertrophic chondrocytes and mature osteocytes, likely due to the difficulties of collecting these matrix-embedding cells during tissue digestion and cell sorting. Thus, we cannot use the sequencing data to compare the Csf1 expression level between those cells and MALPs.

To our surprise, while high trabecular bone mass was evident in long bones, not changes were detected in the vertebrae of *Csf1 CKO*^*Adipoq*^ mice. Based on our scRNA-seq data, Adipoq expression is highly specific for MALPs. Since Csf1 is also expressed at a considerable level in LCPs, we believe that LCP-derived Csf1, which should not be affected in *Csf1 CKO*^*Adipoq*^ mice, might be sufficient for promoting osteoclastogenesis in vertebrae. Future studies identifying specific markers for LCPs and designing LCP-specific *Csf1* knockout mice could add another missing piece of mesenchymal subpopulations that contributes to bone resorption and hematopoiesis.

It is traditionally thought that osteogenic cells, particularly osteocytes, are the major source of RANKL and Csf1 (35). Thus, bone forming cells and bone resorbing cells constitute a feedback loop to maintain the fine balance of bone remodeling. Our findings add MALPs as a new cell type governing this balance. On the one hand, depletion of RANKL or Csf1 in MALPs results in a similar or even more severe osteopetrotic phenotype than depletion of those factors in osteoblasts/osteocytes. In this study, we found that femoral trabecular bone mass increased in *Csf1 CKO*^*Adipoq*^ mice by ∼73% at 3 months of age, comparable to the trabecular bone mass increase in *Csf1 CKO*^*Dmp1*^ mice at 4 months of age (19). In our previous study, we found that *Rankl CKO*^*Adipoq*^ mice develop osteopetrosis at 1 month of age while *Rankl CKO*^*Dmp1*^ mice at the same age have normal bone mass (25). On the other hand, MALPs also suppress bone formation. Using a cell ablation model, we found that depletion of MALPs results in de novo bone formation in diaphyseal bone marrow, which cannot be rescued by transplant of *WT* white adipose tissue (23). Studies by Zuo et al. validated this phenotype and further identified that MALPs highly express two potent BMP signaling inhibitors, chordin-like1 (Chrdl1) and gremlin1 (Grem1), to suppress osteogenic differentiation of bone marrow MSCs (32). By acting on both bone resorption and formation, we believe that MALPs limit the overall trabecular bone mass to provide enough marrow space for blood production. This is also consistent with our recent findings that MALPs highly express hematopoietic factors and that MALPs mediate bone marrow blood cell recovery after radiation damage (36).

Unlike RANKL, which is highly specific for mature osteoclast formation, Csf1 has much broader actions in tissue development, homeostasis, and repair (16). Our previous study with *Rankl CKO*^*Adipoq*^ mice mainly detected bone phenotypes due to suppressed osteoclast formation. We did not notice any changes in bone marrow cellularity in those mice. However, *Csf1 CKO*^*Adipoq*^ mice have hematopoietic phenotypes. Compared to *WT* mice, their bone marrow cellularity, macrophages, and hematopoietic progenitors are reduced. In line with the well-known knowledge that Csf1 drives myelopoiesis from HSCs (28), these results further indicate that MALPs-derived Csf1 plays an important role in hematopoiesis, particularly the production of marrow-resident macrophages. In contrast, a recent study using *Lepr-Cre* to knockout *Csf1* did not find any changes in mouse bone marrow cellularity, marrow blood cells (including HSCs and macrophages), and peripheral blood composition (37). Since MALPs also express *Lepr* (24), we notice the inconsistency between those data and ours. Future experiments should be performed to examine whether *Csf1 CKO*^*Lepr*^ mice have suppressed bone resorption and to compare the *Csf1* expression pattern in bones of *CKO*^*Lepr*^ and *CKO*^*Adipoq*^ mice.

In summary, the present analysis of *Csf1 CKO*^*Adipoq*^ mice contributes significantly to our understanding of cellular sources of Csf1 that regulate bone remodeling and myeloid blood cell production. Together with our previous analysis of *Rankl CKO*^*Adipoq*^ mice, our research demonstrates adipogenic progenitors in the bone marrow are an important mesenchymal subpopulation that controls osteoclastogenesis. The general functions of Csf1 on monocyte lineage cells implicate that MALPs might have broad roles in inflammation and tissue regeneration. Therefore, future directions should be aimed to develop novel therapeutic strategies for bone and blood-related disorders by specifically targeting MALPs.

## Methods

### Analysis of scRNA-seq datasets

Pre-aligned single-cell RNA sequencing (RNA-seq) matrix files were acquired from GEO GSE145477 and GSE176171. Standard Seurat pipeline (38) was used for filtering, normalization, variable gene selection, dimensionality reduction analysis and clustering. For the integrated dataset, batch integration was performed using Harmony (version 1.0) (39). Cell type was annotated according to the metadata from published datasets (23, 29).

### Animals study design

All animal work performed in this report was approved by the Institutional Animal Care and Use Committee (IACUC) at the University of Pennsylvania. *Adipoq-Cre Rosa-tdTomato* (*Adipoq/Td*) mice were generated by breeding *Rosa-tdTomato* (40) mice with *Adipoq-Cre* mice (41). To generate *Csf1 CKO*^*Adipoq*^ mice, we first bred *Adipoq-Cre* (41) with *Csf1*^*flox/flox*^ mice (30) to obtain *Adipoq-Cre Csf1*^*flox/+*^, which were then crossed with *Csf1*^*flox/flox*^ to generate *Csf1 CKO*^*Adipoq*^ mice and *WT* (*Csf1*^*flox/flox*^ and *Csf1*^*flox/+*^) siblings. *Csf1 CKO*^*Prx1*^ mice were generated by a similar breeding strategy with *Prx1-Cre* (42). All mouse lines, except *Csf1*^*flox/flox*^, were obtained from Jackson Laboratory (Bar Harbor, ME, USA). For LPS-induced bone destruction, 6-week-old male mice were injected with 25 mg/kg LPS (Sigma Aldrich, St. Louis, MO) or PBS above calvariae. After 7 days, calvariae were collected and analyzed by microCT followed by TRAP staining.

### Micro-computed tomography (microCT) analysis

MicroCT analysis (microCT 35, Scanco Medical AG, Brüttisellen, Switzerland) was performed at 6 µm isotropic voxel size as described previously (43). Briefly, the distal end of femur corresponding to a 0 to 2.8 mm region below the growth plate was scanned at 6 μm isotropic voxel size to acquire a total of 466 μCT slices per scan. The images of the secondary spongiosa regions (0.6 to 1.8 mm below the lowest point of the growth plate) were contoured for trabecular bone analysis. At the femur midshaft, a total of 100 slices located 4.8-5.4 mm away from the distal growth plate were acquired for cortical bone analyses by visually drawing the volume of interest (VOI). In vertebrae, the region 50 slices away from the top and bottom end plates (∼300 slices) was acquired for trabecular bone analysis. The trabecular bone tissue within the VOI was segmented from soft tissue using a threshold of 487.0 mgHA/cm^3^ and a Gaussian noise filter (sigma=1.2, support=2.0). The cortical bone tissue was segmented using a threshold of 661.6 mgHA/cm^3^ and a Gaussian noise filter (sigma=1.2, support=2.0). Three-dimensional standard microstructural analysis was performed to determine the geometric trabecular bone volume fraction (BV/TV), bone mineral density (BMD), trabecular thickness (Tb.Th), trabecular separation (Tb.Sp), trabecular number (Tb.N), and structure model index (SMI). For analysis of cortical bone, periosteal perimeter (Ps.Pm), endosteal perimeter (Ec.Pm), cortical bone area (Ct.Ar), cortical thickness (Ct.Th), and tissue mineral density (TMD) were recorded. All calculations were performed based on 3D standard microstructural analysis (44).

Calvariae were scanned at 15 µm isotropic voxel size. The three-dimensional images were reconstructed to visualize the destructive area. A square region of 8 mm x 8 mm centered at the midline suture was selected for further quantitative analysis by ImageJ.

### Histology and bone histomorphometry

To obtain whole mount sections for immunofluorescent imaging of *Csf1 CKO*^*Adipoq*^ mouse bones, freshly dissected femurs were fixed in 4% PFA for 1 day, decalcified in 10% EDTA for 4-5 days, and then immersed into 20% sucrose and 2% polyvinylpyrrolidone (PVP) at 4°C overnight. Then samples were embedded into medium containing 8% gelatin, 20% sucrose and 2% PVP and sectioned at 50 µm in thickness. Sections were incubated with rat anti-Endomucin (Santa cruz, sc-65495),), rabbit anti-osterix (Abcam, ab22552), rat anti-F4/80 (Biolegend, 123101), or rabbit anti-Perilipin (Cell signaling, 9349) at 4°C overnight followed by Alexa Fluor 488 anti-rat (Abcam, ab150155) or Alexa Fluor 647 anti-rabbit (Abcam, ab150075) secondary antibodies incubation 1 hour at RT.

To obtain paraffin sections, mouse calvaria bones were fixed in 4% PFA for 24 hr and decalcified in a 10% EDTA for 5-7 days at 4°C. Samples were then embedded in paraffin, sectioned at 6 μm in thickness, and processed for tartrate-resistant acid phosphatase (TRAP) staining using Acid Phosphatase, Leukocyte (TRAP) Kit (Sigma-Aldrich, 387A) with fast green counterstaining.

To obtain cryosections, mouse bones were dissected and fixed in 4% PFA for 24 hr, dehydrated in 30% sucrose in PBS, embedded in optimal cutting temperature (OCT) compound, and sectioned at 6 μm in thickness using a cryofilm tape (Section Lab, Hiroshima, Japan). Fluorescent TRAP staining was performed as described previously (45). For dynamic histomorphometry, mice received calcein (10 mg/kg, Sigma Aldrich) and xylenol orange (90 mg/kg, Sigma Aldrich) at 9 and 2 days, respectively, before euthanization. Sagittal cryosections of tibiae prepared with cryofilm tape were used for dynamic histomorphometry. Sections were scanned by a Nikon Eclipse 90i fluorescence microscope and areas within secondary spongiosa were quantified by OsteoMeasure Software (OsterMetrics, Decatur, GA, USA). The primary indices include total tissue area (TV), trabecular bone perimeter (BS), single- and double-labeled surface, and interlabel width. Mineralizing surface (MS) and surface-referent bone formation rate (BFR/BS, μm^3^/μm^2^/d) were calculated as described by Dempster et al. (46).

### Hematopoietic phenotyping of bone marrow cells

Bone marrow was flushed from femurs of 3-month-old *WT* and *Csf1 CKO*^*Adipoq*^ mice. The lineage cell compartment of the bone marrow was analyzed by staining for myeloid (rat anti-mouse Gr-1 APC-Cy7, BD, 557661, rat anti-mouse Mac-1 APC, eBioscience, 17-0112-83), lymphoid lineages (rat anti-mouse B220 FITC, eBioscience, 11-0452-82, hamster anti-mouse CD3 PE-Cy7, eBioscience, 25-0031-82), erythroid lineages (rat anti-mouse Ter119 APC, BD, 557909) or megakaryocytes (rat anti-mouse CD41 FITC). The HSPC compartment was analyzed by staining for Lineage markers biotin-Ter-119, -Mac-1, -Gr-1, -CD4, -CD8α, -CD5, -CD19 and -B220 (eBioscience, 13-5921-85, 13-0051-85, 13-5931-86, 13-0112-86, 13-0452-86, 13-0041-86, 13-0081-86, 13-0193-86) followed by staining with streptavidin-PE-Texas Red (Invitrogen, SA1017), rat anti-mouse cKit APC-Cy7 (eBioscience, 47-1171-82), rat anti-mouse Sca1 PerCP- Cy5.5 (eBioscience, 45-5981-82), hamster anti-mouse CD48 APC (eBioscience, 17-0481-82) and rat anti-mouse CD150 PE-Cy7 (Biolegend, 115914). To examine macrophages, flushed bone marrow cells were stained with rat anti-mouse F4/80 BV711 (Biolegend, 123147) and rat anti-mouse CD11b FITC (Biolegend, 101205). All flow cytometry experiments were performed by either a LSR A or a BD LSR Fortessa flow cytometer and analyzed by FlowJo v10.5.3 for MAC. *Cell culture*

For in vitro osteoclastogenesis, bone marrow cells flushed from long bones were seeded to dishes for obtaining bone marrow macrophages (BMMs) as described previously (47, 48). BMMs were then seeded at 1 × 10^6^ cells/well in 24-well plates and stimulated with 30 ng/mL RANKL and 30 ng/mL M-CSF (R&D Systems, Minneapolis, MN, USA) for 5 days to generate mature OCs. TRAP staining was performed using a TRAP kit (Sigma Aldrich, 387A). Osteoclasts were quantified by counting the number of TRAP^+^, multinucleated cells (>3nuclei/cell) per well.

### Serum biochemistry assay

Sera were collected from *WT* and *Csf1 CKO*^*Adipoq*^ mice during euthanization for measuring bone turnover markers, collagen type I C-telopeptide degradation products (mouse CTX-I Elisa kit; AFG Scientific, Northbrook, IL, USA) and N-terminal propeptide of type I procollagen (Immunotag(tm) Mouse PINP ELISA Kit, G-Bioscience, St. Louis, MO, USA) respectively, according to the manufacturer’s instructions.

### RNA analysis

Sorted cells or bone marrow tissue was collected in Tri Reagent (Sigma Aldrich) for RNA purification. A Taqman Reverse Transcription Kit (Applied BioSystems, Inc., Foster City, CA, USA) was used to reverse transcribe mRNA into cDNA. The power SYBR Green PCR Master Mix Kit (Applied BioSystems, Inc) was used for quantitative real-time PCR (qRT-PCR). Primers for *Csf1* gene are 5’-ATGAGCAGGAGTATTGCCAAGG-3’ (forward) and 5’- TCCATTCCCAATCATGTGGCTA-3’ (reverse), and primes for β-actin gene are 5’- GGCTGTATTCCCCTCCATCG-3’(forward) and 5’-CCAGTTGGTAACAATGCCATGT-3’ (reverse).

### Statistical analyses

Data are expressed as means ± standard deviation (SD). For comparisons between two groups, unpaired two-sample student’s t-test was applied. For comparisons amongst multiple groups across two fixed effect factors (e.g., genotype and age), two-way ANOVA was applied, followed by Tukey-Kramer multiple comparison test to account for family-wise type I error using Prism 8 software (GraphPad Software, San Diego, CA). In all tests, the significance level was set at α = 0.05. For assays using primary cells, experiments were repeated independently at least three times and representative data were shown here. Values of p<0.05 were considered statistically significant.

## Acknowledgments

We thank the MicroCT Imaging Core at Penn Center for Musculoskeletal Disorders (PCMD) for their assistance with microCT scanning and analysis. This study was supported by NIH grants NIA R01AG069401, NIAMS R21AR078650 (to L.Q.), R00AR067283 (to N.D.), P30AR069619 (to PCMD), R01AG045040 (to J.X.J) and Welch Foundation grant (AQ-1507) (to JXJ).

## Supplementary Materials

**Figure S1.**
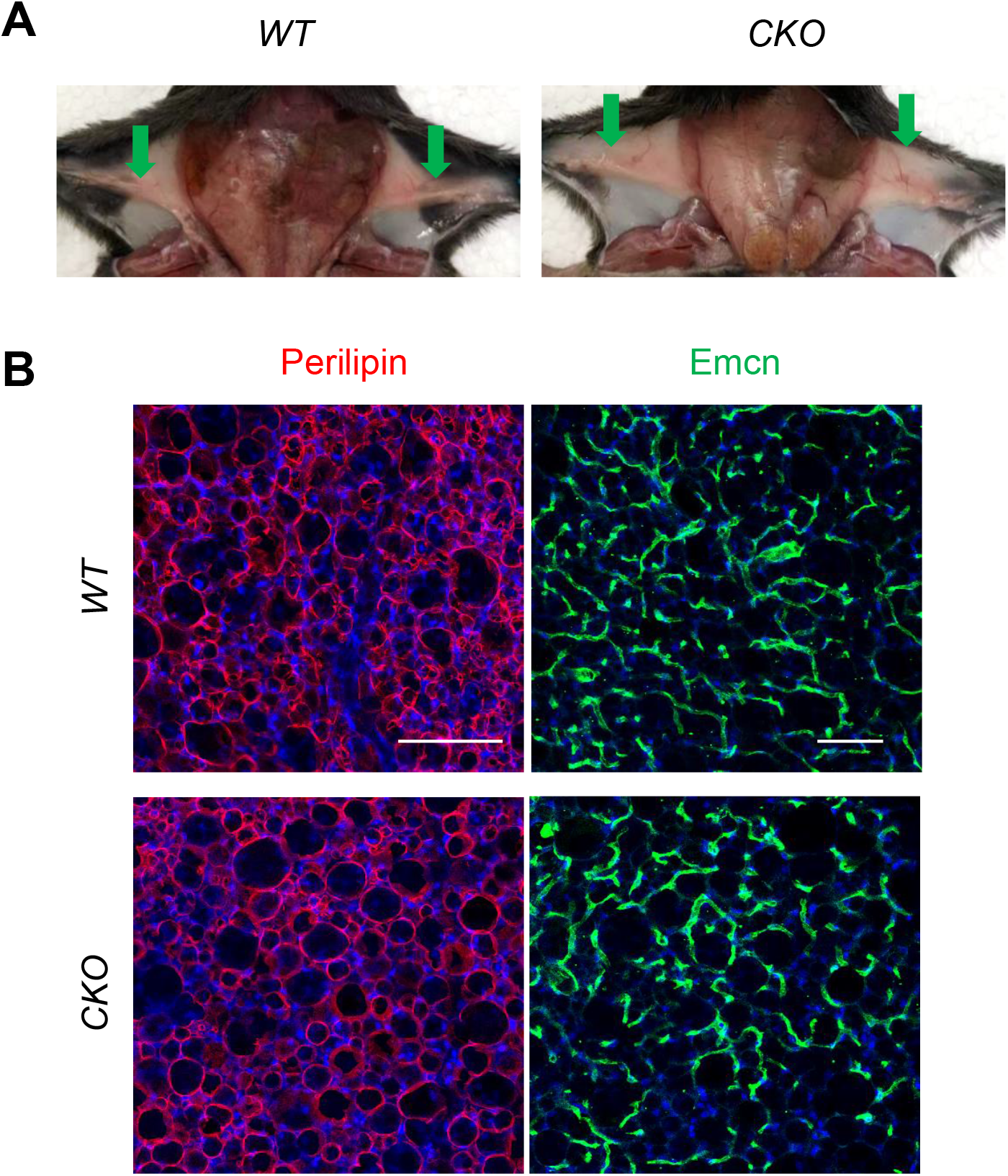
*Csf1 CKO*^*Adipoq*^ mice have normal subcutaneous fat pad. (A) Representative photographs showing subcutaneous fat pad tissues (pointed by arrows) in *WT* and *Csf1 CKO*^*Adipoq*^ mice. (B) Representative adipocyte (Perilipin) and vessel (Emcn) staining images in subcutaneous fat. Scale bar=100 µm (Perilipin) or 200 µm (Emcn).

**Figure S2.**
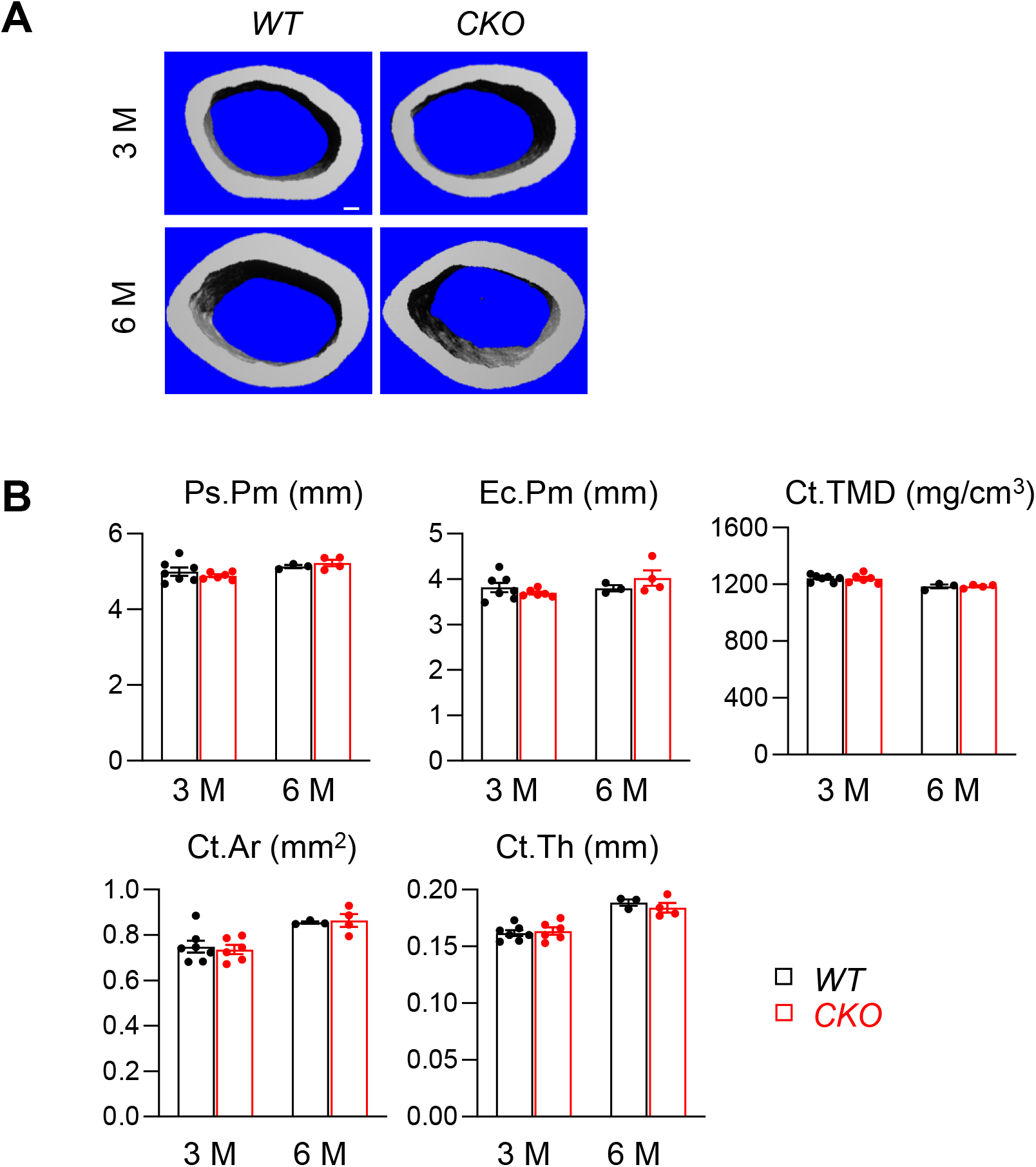
*Csf1 CKO*^*Adipoq*^ mice have normal cortical bone structure. (A) 3D microCT reconstructions of the femoral midshaft region from 3 and 6 month-old female *WT* and *Csf1 CKO*^*Adipoq*^ mice. Scale bar=100 µm. (B) MicroCT measurement of cortical bone structural parameters. Ps.Pm: periosteal perimeter; Ec.Pm: endosteal perimeter; Ct.TMD: cortical tissue mineral density. Ct.Ar: cortical area; Ct.Th: cortical thickness. n=3-7 mice/group.

**Figure S3.**
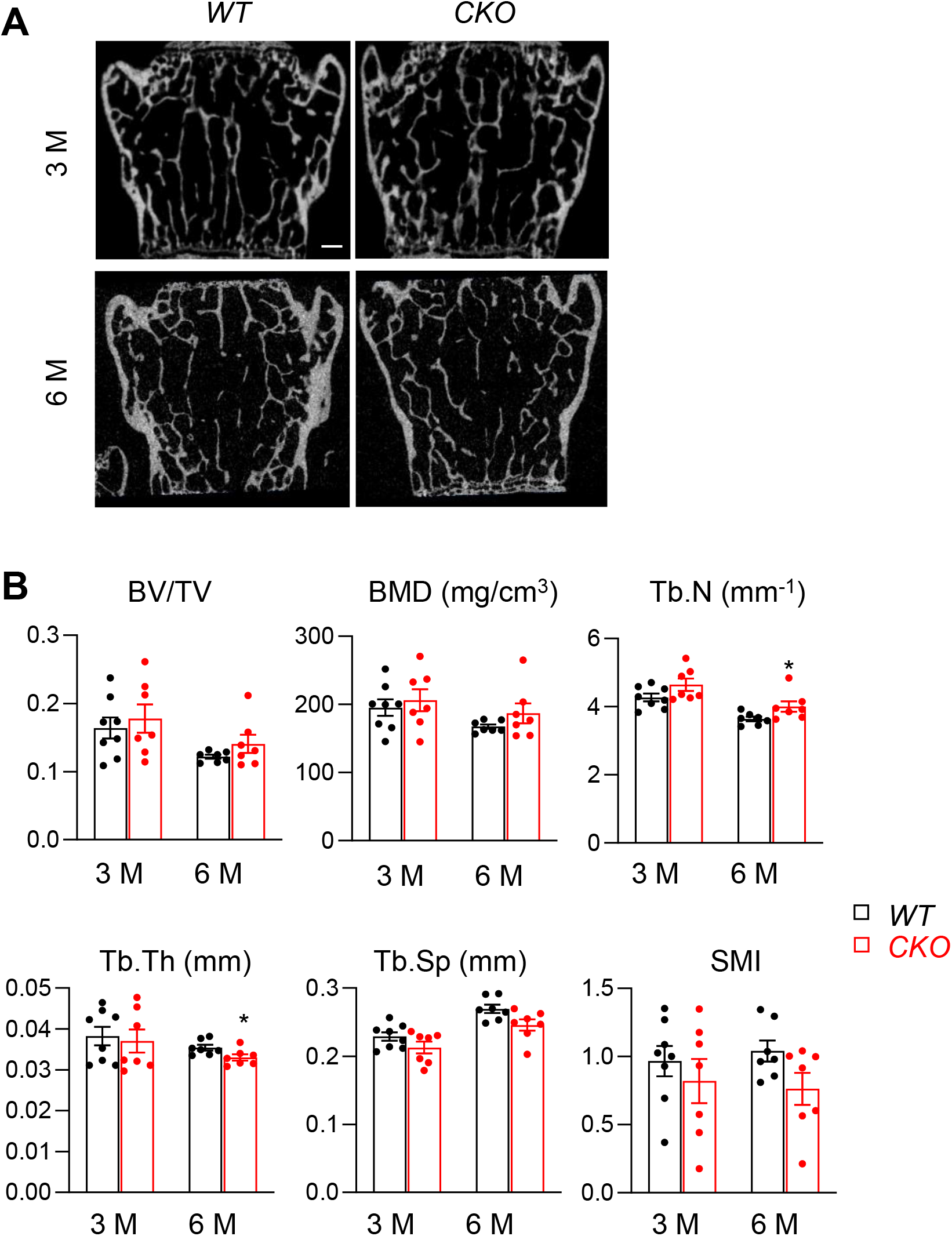
*Csf1* deficiency in MALPs does not affects vertebral bone. (A) 2D microCT reconstructions of vertebrae from *WT* and *Csf1 CKO*^*Adipoq*^ mice at 3- and 6- months. Scale bar=250 µm. (B) MicroCT measurement of trabecular bone structural parameters in vertebrae. BV/TV: bone volume fraction; BMD: bone mineral density; Tb.N: trabecular number; Tb.Th: trabecular thickness; Tb.Sp: trabecular separation; SMI: structural model index. n=6–7 mice/group. *, p<0.05 *CKO* vs *WT*.

**Figure S4.**
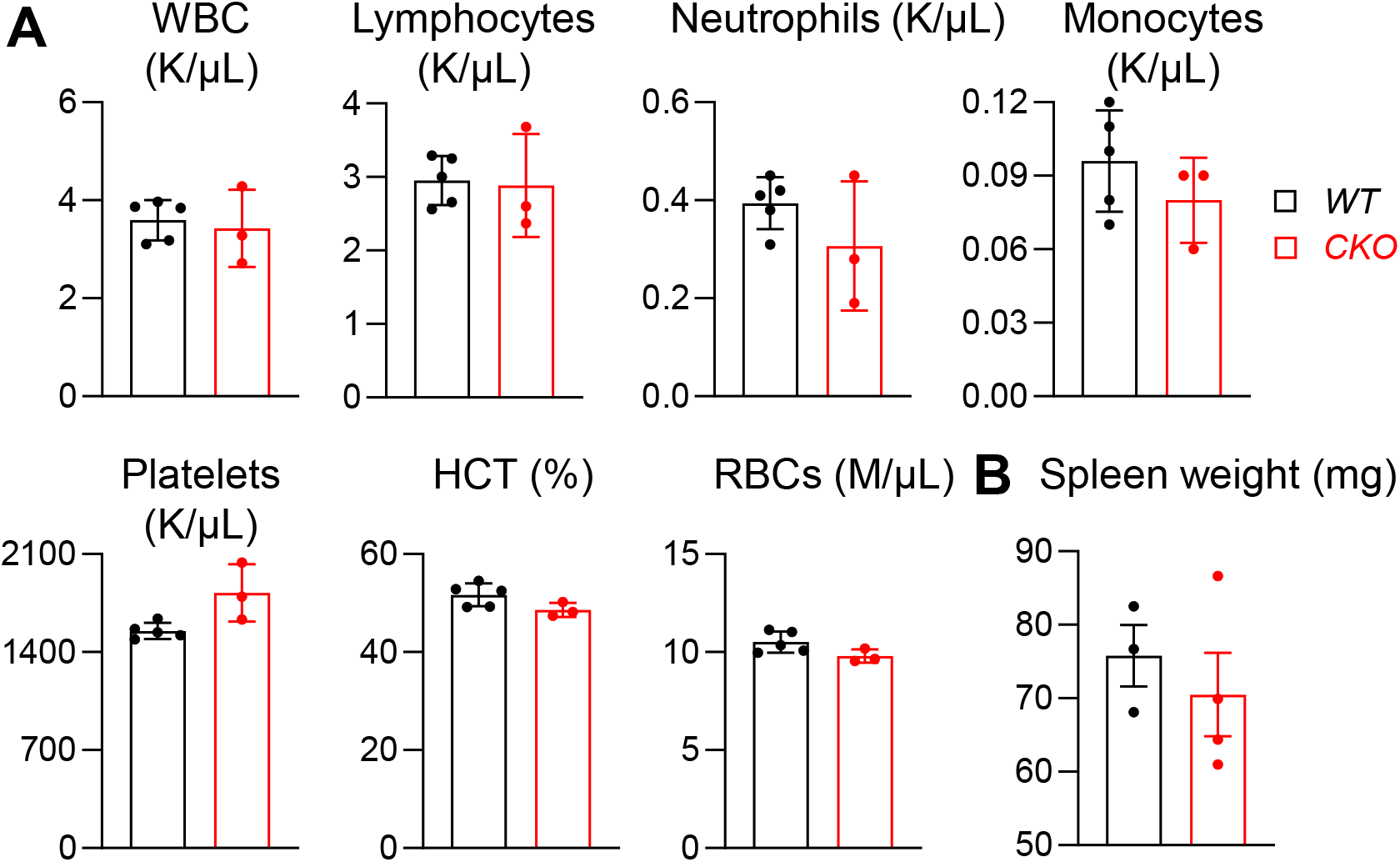
Peripheral blood and spleen appear normal in *Csf1 CKO*^*Adipoq*^ mice. (A) Flow analysis of peripheral blood components, including white blood cells (WBCs), lymphocytes, neutrophils, monocytes, platelets, hematocrit (HCT), and red blood cells (RBCs) in *WT* and *Csf1 CKO*^*Adipoq*^ mice at 3 months of age. n=3-5 mice/group. (B) Spleen weight is measured at 3 months of age. n=3-4 mice/group.

